# Elevated Neuropeptide Y1 Receptor Signaling Contributes to β-cell Dysfunction and Failure in Type 2 Diabetes

**DOI:** 10.1101/2021.05.14.444149

**Authors:** Chieh-Hsin Yang, Danise Ann-Onda, Xuzhu Lin, Stacey Fynch, Shaktypreya Nadarajah, Evan Pappas, Xin Liu, John W. Scott, Jonathan S. Oakhill, Sandra Galic, Yanchuan Shi, Alba Moreno-Asso, Cassandra Smith, Tom Loudovaris, Itamar Levinger, Decio L. Eizirik, Ross D. Laybutt, Herbert Herzog, Helen E. Thomas, Kim Loh

**Affiliations:** St. Vincent’s Institute of Medical Research, Fitzroy, VIC, 3065, Australia; Mary MacKillop Institute for Health Research, Australian Catholic University, Melbourne VIC, 3000, Australia; The Florey Institute of Neuroscience and Mental Health, Parkville, VIC, 3052, Australia; Department of Medicine, University of Melbourne, Fitzroy, VIC, 3065, Australia; Garvan Institute of Medical Research, St Vincent’s Hospital, Sydney, 2010, Australia.; Faculty of Medicine, UNSW Australia, Sydney, 2052, Australia.; Institute of Health and Sport (IHES), Victoria University, Footscray, VIC, Australia; Australian Institute for Musculoskeletal Science (AIMSS), University of Melbourne and Western Health, St Albans, VIC Australia; ULB Center for Diabetes Research, Medical Faculty, Universite Libre de Bruxelles (ULB), Brussels, Belgium; Indiana Biosciences Research Institute (IBRI), Indianapolis, Indiana, USA

**Author notes:** Corresponding authors: Kim Loh, PhD Head, Diabetes and Metabolic Disease laboratory St. Vincent’s Institute of Medical Research 9 Princes Street Fitzroy, VIC, 3065, Australia Chieh-Hsin Yang, PhD St. Vincent’s Institute of Medical Research 9 Princes Street Fitzroy, VIC, 3065, Australia.

**Keywords:** NPY, β-cell, insulin secretion, Y1 receptor, type 2 diabetes

## Abstract

Loss of functional β-cell mass is a key factor contributing to the poor glycaemic control in type 2 diabetes. However, therapies that directly target these underlying processes remains lacking. Here we demonstrate that gene expression of neuropeptide Y1 receptor and its ligand, neuropeptide Y, was significantly upregulated in human islets from subjects with type 2 diabetes. Importantly, the reduced insulin secretion in type 2 diabetes was associated with increased neuropeptide Y and Y1 receptor expression in human islets. Consistently, pharmacological inhibition of Y1 receptors by BIBO3304 significantly protected β-cells from dysfunction and death under multiple diabetogenic conditions in islets. In a preclinical study, Y1 receptor antagonist BIBO3304 treatment improved β-cell function and preserved functional β-cell mass, thereby resulting in better glycaemic control in both high-fat-diet/multiple low-dose streptozotocin- and *db/db* type 2 diabetic mice. Collectively, our results uncovered a novel causal link of increased islet NPY-Y1 receptor signaling to β-cell dysfunction and failure in human type 2 diabetes. These results further demonstrate that inhibition of Y1 receptor by BIBO3304 represents a novel and effective β-cell protective therapy for improving functional β-cell mass and glycaemic control in type 2 diabetes.

## INTRODUCTION

The prevalence of diabetes has been increasing over the last few decades and is now a major health concern worldwide [1]. It is a major cause of premature mortality and other health complications such as cardiovascular disease and chronic kidney disease [2, 3]. Located within the islets of Langerhans, pancreatic β-cells synthesize the hormone insulin, which is secreted primarily in response to elevated blood glucose levels. Type 2 diabetes (T2D) is the result of insufficient production of the glucose-lowering hormone insulin [4, 5], triggered by multiple factors. Peripheral insulin resistance, coupled with diabetogenic stressors including hyperlipidaemia, endoplasmic reticulum (ER), oxidative stresses and inflammation are recognised as major driving forces of β-cell dysfunction and death, which ultimately leads to or exacerbates hyperglycaemia, a key hallmark of T2D [6–8]. Current therapies for the treatment of T2D are mainly focused on improving glycaemic control through increased insulin secretion from the β-cell and/or the improvement of insulin sensitivity. While T2D management has improved over the years, the search for a novel agent that can selectively improve β-cell function together with the preservation of β-cell mass remains ongoing.

The neuropeptide Y system consists of neuropeptide Y (NPY), peptide-YY (PYY) and pancreatic polypeptide (PP), are a group of short (36-amino acid) peptides that play a key role in the regulation of energy homeostasis [9, 10]. While NPY centrally promotes feeding and reduces energy expenditure, PYY and PP mediate satiety [9, 10]. The NPY system exerts its biological actions via a set of G-protein-coupled receptors (GPCR), of which five have been cloned: Y1, Y2, Y4, Y5, and y6 [9, 10]. The NPY system is widely expressed in the central nervous system as well as in peripheral tissues [9, 10]. In the pancreas, while PYY and PP are expressed by α-cells and pancreatic PP cells, respectively, recent studies have demonstrated that NPY expression in mouse islet β-cells may play a role in altered β-cell function that precedes to diabetes onset [10, 11]. Interestingly, NPY levels were significantly upregulated in response to oxidative stress in islets from subjects with T2D [11]. Furthermore, NPY-deficient mice exhibit enhanced insulin secretion in response to glucose administration [12]. This is further confirmed by *in vitro* studies demonstrating that application of NPY decreases glucose-stimulated insulin secretion from mouse islets [12]. Together, these results suggest that NPY may act through a paracrine mechanism to tonically suppress β-cell function.

In addition to the brain, we previously identified that neuropeptide Y1 receptor is also expressed in mouse and human β-cells and acts as a critical negative regulator of β-cell function [10, 13, 14]. Like all Y-receptors, the Y1 receptor is a GPCR that preferentially associates with Gi/o G-protein and therefore acts in an inhibitory fashion reducing cyclic AMP (cAMP) levels. Indeed, we have shown that pharmacological inhibition of this receptor using a Y1 receptor specific antagonist, BIBO3304, significantly enhances β-cell function via a cAMP-dependent mechanism in mouse and human islets [13]. In addition, we demonstrated that BIBO3304 delays the onset of T1D and may also be useful in boosting β-cell function under conditions where insulin secretion is limited such as during islet transplantation [13]. However, the beneficial effects of pharmacological inhibition of the Y1 receptor in T2D remain unknown. Here, we show in proof-of-concept studies that Y1 receptor antagonist BIBO3304 acts as a β-cell protective agent. BIBO3304 treatment significantly improved glycaemic control in the high-fat diet/multiple low-doses streptozotocin-induced and obese leptin receptor deficient (*db/db*) T2D mouse models. Importantly, our findings have direct relevance to the clinical setting of T2D, since BIBO3304 exhibited equal efficacy in improving glycaemic control as the first line oral anti-diabetic drug, metformin.

## RESULTS

### Increased NPY and Y1 receptor levels in T2D islets are associated with reduced insulin secretion

To investigate whether the NPY system in pancreatic islets is associated with reduced β-cell function in the pathogenesis of T2D, we first determined the NPY system expression profiles in human islets isolated from non-diabetic and T2D subjects as described **Methods and Supplementary Data.** Interestingly, we found that in islets from diabetic donors, the expression of *NPY* and its receptor *NPY1R* was increased by 2.7- and 2.5-fold, respectively, as compared to the non-diabetic donors (Figure 1A and 1B). Importantly, the increased *NPY* and *NPY1R* mRNA expression in human islets correlated with reduced insulin secretion as indicated by the insulin stimulation index (Spearman’s r=0.7151, p=0.0376 and r=0.6524, p=0.0473) (Figure 1C and 1E), whereas the differential expression of *NPY* and *NPY1R* was not associated with HbA1c (Figure 1D and 1F) or BMI (Figure S1A and S1B). These results suggests that elevated NPY/Y1 receptor signaling may contribute to impaired insulin secretion in human with T2D.

**Figure 1.**
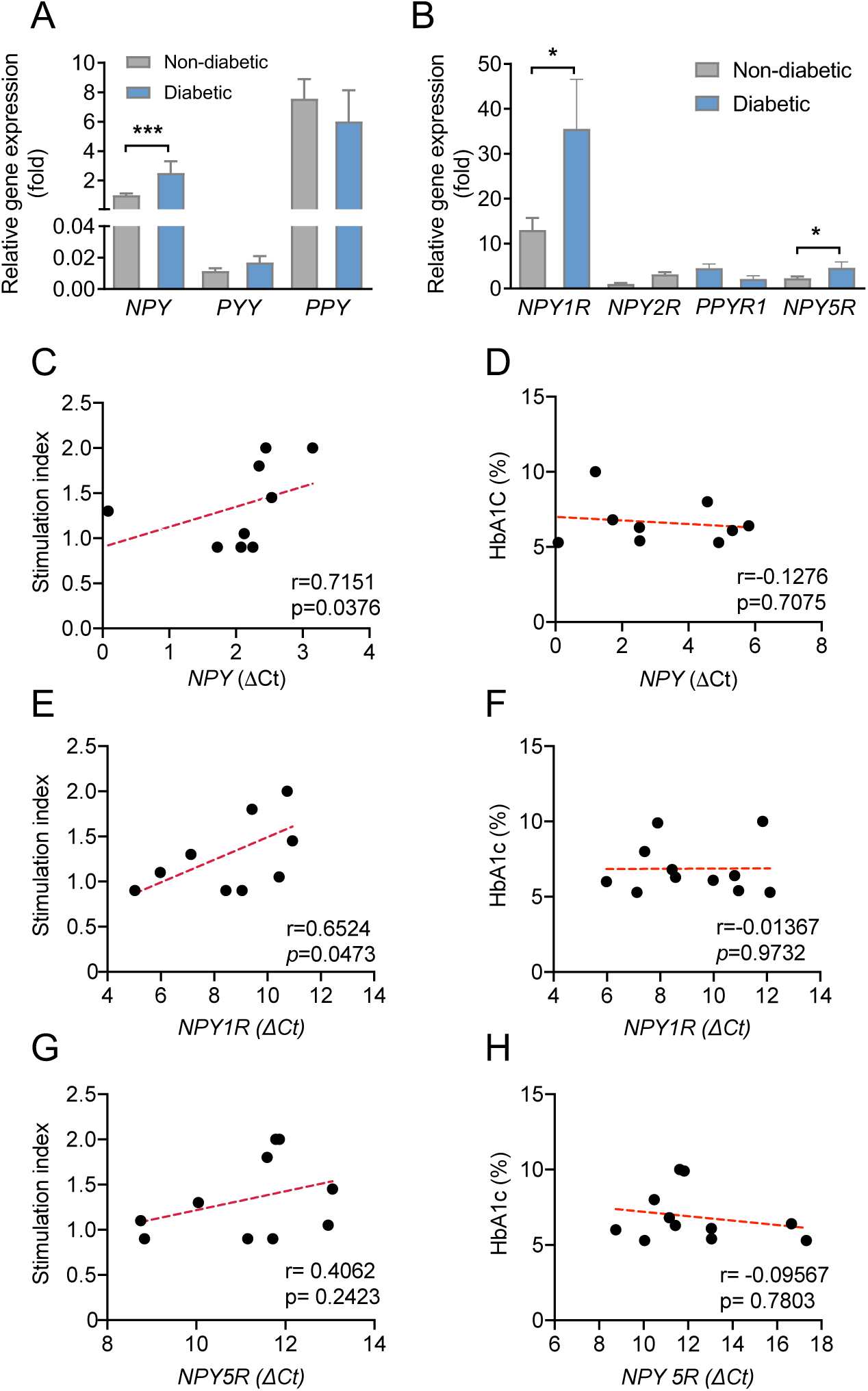
Increased NPY and Y1 receptor mRNA expression levels negatively correlate with islet stimulation index in T2D. (A) *NPY*, *PYY* and *PPY* mRNA expression in human pancreatic islets from non-diabetic and T2D subjects relative to the *NPY* expression in non-diabetic group. Subject numbers: non-diabetic = 25 and type 2 diabetic = 11. (B) Y-receptor expression profiles in human pancreatic islets from non-diabetic and T2D subjects relative to the *NPY2R* expression in the non-diabetic group. Subject numbers: non-diabetic = 25 and type 2 diabetic =11. (C-D) Correlation between the insulin stimulation index or HbA1C and the expression of *NPY* mRNA (delta CT values) in human islets of subjects with T2D and non-diabetic control subjects. Total subjects = 9. (E-F) Correlation between the insulin stimulation index or HbA1C and the expression of *NPY1R* mRNA (delta CT values) in human islets of subjects with T2D and non-diabetic control subjects. Total subjects = 10. (G-H) Correlation between the insulin stimulation index or HbA1C and the expression of *NPY5R* mRNA (delta CT values) in human islets of subjects with T2D and non-diabetic control subjects. Total subjects = 11. (A-B) Data are mean ± SEM. *P* values by two-tailed t-test when comparing non-diabetic vs diabetic. (C-H) *P* values by two-tailed Spearman correlation analysis.

On the other hand, the basal levels of *NPY2R*, *PPYR1* (also known as *NPY4R*) and *NPY5R* were very low in pancreatic islets (Figure 1B), suggesting that these Y receptors are unlikely to play a major role in mediating NPY function in human pancreatic islets. Despite the low level of expression, *NPY5R* was also moderately upregulated in islets of T2D subjects (Figure 1B), but there was no significant correlation between *NPY5R* expression and insulin stimulation index, HbA1c (Figure 1G and 1H) or BMI (Figure S1C). Importantly, the NPY/Y1 receptor axis appeared to be exclusively up-regulated as there were no noticeable changes in other NPY ligands such as *PYY* and *PPY* and their correlation with insulin stimulation index or HbA1c (Figure S1D-1I). Collectively, these results uncover a novel link between increased islet NPY/Y1 receptor signaling and β-cell dysfunction in human T2D.

### Y1 receptor inhibition restores β-cell function and protects against β-cells apoptosis under diabetogenic conditions

Diabetogenic stresses such as inflammation, ER stress, oxidative stress and glucolipotoxicity have been implicated as key factors contributing to impaired β-cell function and death in T2D [15–17]. Given that islet NPY/Y1 receptor levels negatively correlate with insulin stimulation index in human T2D, we next asked whether pharmacological inhibition of NPY/Y1 receptor signalling by a selective Y1 receptor antagonist BIBO3304, under diabetogenic conditions, would restore β-cell function. To test this, we assessed glucose-stimulated insulin secretion (GSIS) with or without BIBO3304 on wild-type C57BL/6 islets that were exposed to various stress conditions (inflammation: proinflammatory cytokines TNFα, IFNγ and IL1β; ER stress: thapsigargin; oxidative stress: H_2_O_2_; glucolipotoxicity: high glucose/palmitate). A significant reduction in insulin release in response to glucose stimulation was observed in islets that were exposed to inflammatory cytokines (Figure 2A), ER-stress (Figure 2B) or H_2_O_2_ (Figure S2A). In contrast, BIBO3304 co-treatment prevented the impaired insulin release that was induced by proinflammatory cytokines or ER stress inducer, thapsigargin (Figure 2A and 2B) with the same trend observed in high glucose/palmitate and oxidative stress (Figure S2A) treated islets. Taken together, these results demonstrate that, under diabetogenic stress conditions such as inflammation and ER stress, Y1 receptor antagonism can directly restore β-cell function by enhancing GSIS.

**Figure 2.**
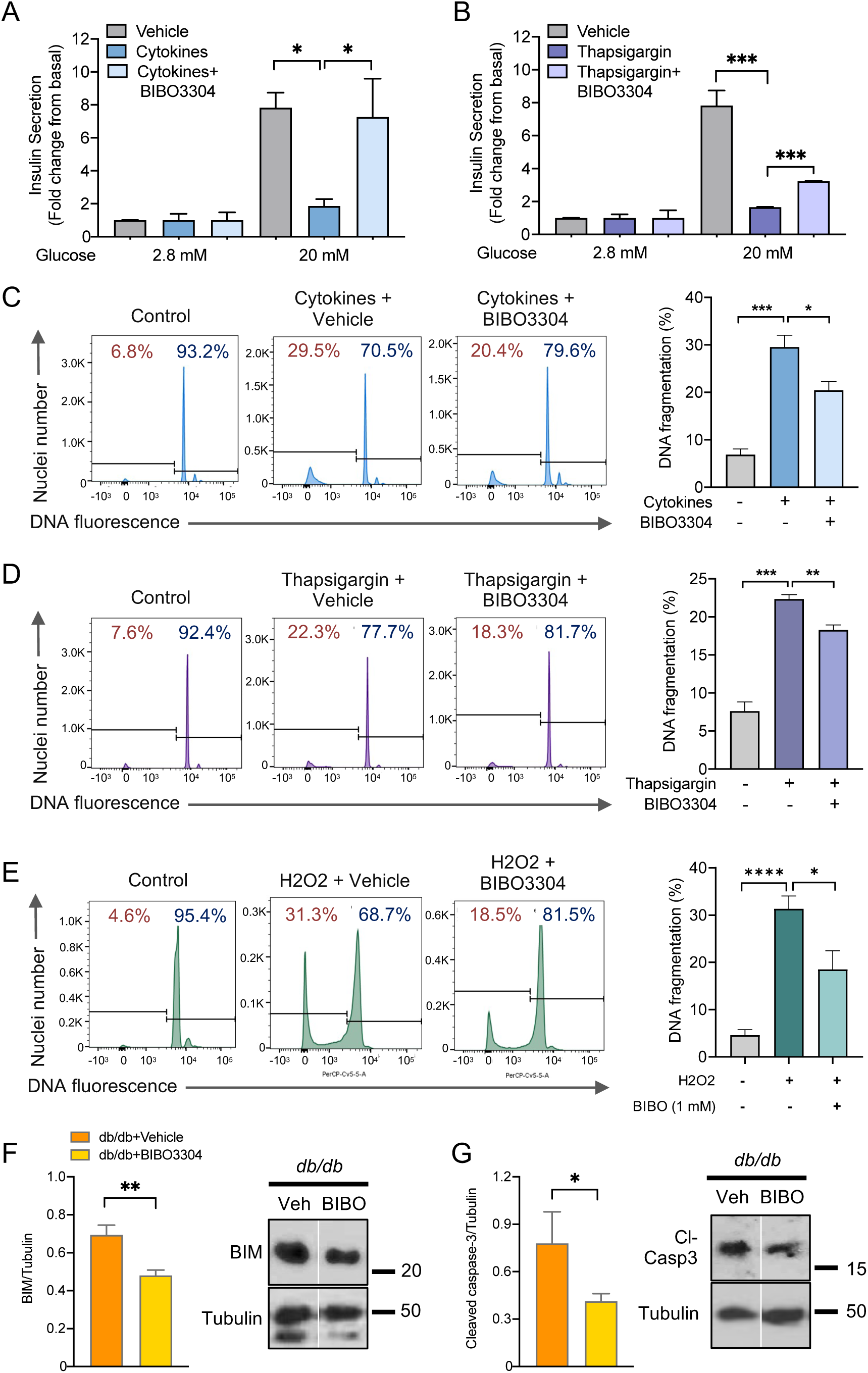
Pharmacological inhibition of Y1 receptor restores β-cell function and protects against β-cell death under diabetogenic conditions. (A-B) Pancreatic islets from C57BL/6 mice were isolated and cultured in the corresponding diabetogenic conditions: inflammatory cytokine cocktail of 25 ng/ml IL-1β, 250 ng/ml IFNγ, 50 ng/ml TNFα ± 1 μM of BIBO3304 for 48h (*n* = 5), thapsigargin (1 μM) ± 1 μM of BIBO3304 for 24h (*n* = 3-6) (*n* = 3). Glucose-stimulated insulin secretion was determined in response to 2.8 and 20 mmol/L glucose. (C-E) DNA fragmentation in response to inflammation, ER stress and oxidative stress was measured by flow cytometry. Representative FACS profiles are shown and the results are representative of islets from a minimum of 3 mice per group. (F-G) Western blot analyses of pro-apoptotic proteins BIM and cleaved caspase-3 in isolated islets from 10-week-old leptin receptor-deficient *db/db* mice were cultured with/without 1 μM of BIBO3304 for 36h. α-tubulin was used as the loading control (*n* = 3-4). Results shown are a representative blot and quantitative densitometry analysis. Data are mean ± SEM. \**P* < 0.05, \*\**P* < 0.01, calculated by unpaired Student’s *t*-test.

Targeting β-cell preservation is a key component of therapeutic strategies for glycaemic control in diabetes [18]. We next investigated whether inhibition of Y1 receptor could also protect β-cells from failure under diabetogenic conditions by exposing islets to various stress conditions with or without the treatment of Y1 receptor antagonist BIBO3304. Cell death was measured using propidium iodide staining and analysis of the sub-diploid DNA content by flow cytometry. As expected, chronic exposure to all diabetogenic stressors induced a substantial increase in β-cell apoptosis (Figure 2C, 2D **and** 2E). In contrast, BIBO3304 significantly reduced cytokine, thapsigargin and H_2_O_2_-induced cell death (Figure 2C, 2D **and** 2E), but not glucolipotoxicity-induced cell death (Figure S2B), as indicated by decreased subdiploid DNA content. Given that the pathogenesis of T2D is the result of complex metabolic perturbations, islets cultured with individual diabetogenic stress alone may have a limited ability to model the islet microenvironment in T2D. We therefore tested the ability of BIBO3304 to protect β-cells from death in islets isolated from severely diabetic *db/db* mice (random blood glucose (RBG) 27.5±1.84 mmol/L; body weight (BW) 42.4±1.56 g, n=13) compared to their non-diabetic littermate control *db/+* mice (RBG 10.5±0.4 mmol/L; BW 24.5±1.16 g, n=9). Our results showed that the addition of BIBO3304 for 36 h alleviated β-cell apoptosis in *db/db* islets as evidenced by significant reduction in the expression of pro-apoptotic markers such as BIM and cleaved-Caspase-3 (Figure 2F and 2G). Taken together, these results suggest that the direct inhibition of Y1 receptor signalling in islets was associated with reduced apoptosis, indicating that Y1 receptor antagonism plays a protective role in preserving β-cell mass and function in T2D.

### Inhibition of Y1 receptor restores normoglycaemia in high fat diet (HFD) and streptozotocin (STZ)-induced T2D mouse model

While pharmacological inhibition of Y1 receptor restored β-cell function and decreased β-cell death *ex vivo*, we next investigated whether BIBO3304 could improve glycaemic control in the context of a non-genetic animal model of human adult-onset T2D that displays hyperglycaemia and β-cell dysfunction. To test this, C57BL/6 mice were rendered diabetic by HFD diet feeding for 4 weeks and followed by injection with multiple low-dose streptozotocin (STZ) to induce partial β-cell loss. Diabetic HFD/STZ mice were subsequently treated with placebo or BIBO3304 for up to 6 weeks to assess its ability to restore normoglycemia (Figure 3A). Significant hyperglycaemia (blood glucose >15 mmol/L) established 7 days after STZ treatment (Figure 3B). Body weight, adiposity and food intake were comparable between BIBO3304 and placebo treated group (Figure S3A-S3H). Importantly, however, BIBO3304 treated mice displayed significantly lower blood glucose levels during the entire 4 weeks of the study when compared to the placebo group (Figure 3B). Fed and fasted blood glucose levels were also significantly reduced in BIBO3304-treated mice (Figure 3C). The reduced blood glucose levels were unlikely due to an increase in urinary glucose excretion as urine glucose was 2-3 folds lower in BIBO3304 treated mice than in placebo group (Figure 3D). To compare the effects of BIBO3304 with a currently available oral anti-diabetic drug, we tested the effects of metformin, the first-line and most widely prescribed drug for the treatment of T2D. Metformin treatment resulted in improved blood glucose levels in HFD/STZ diabetic mice (Figure 3E) and was not significantly different to the improvement in glycaemic control by BIBO3304 (Figure 3E).

**Figure 3.**
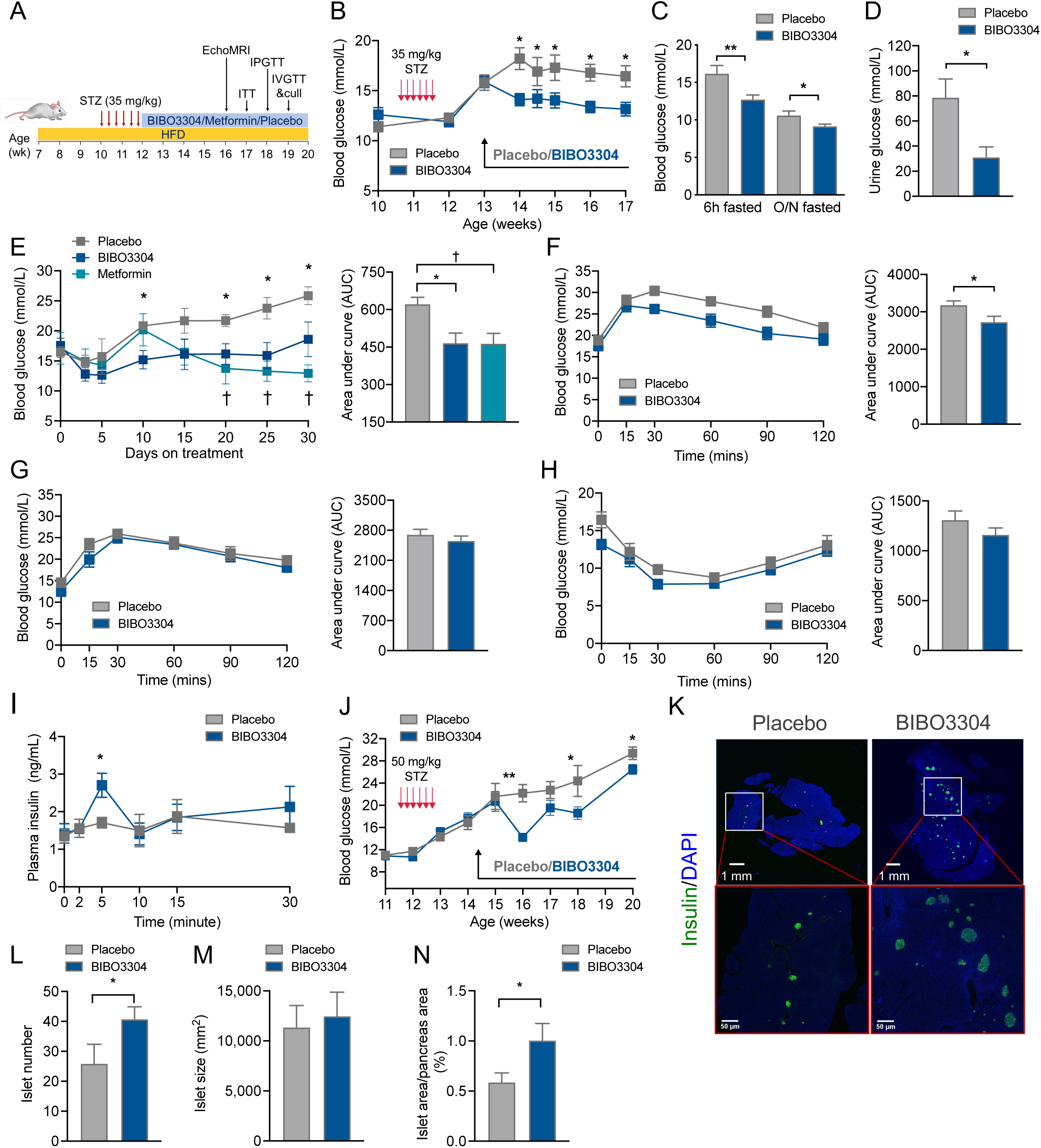
Y1 receptor antagonist BIBO3304 improves glycemia in HFD/STZ-induced diabetes mice. (A) Schematic diagram of treatment regimen. C57BL/6 mice were fed on a high fat diet for 4 weeks and rendered diabetic by multiple low-dose of STZ injections (6 doses, 35mg/kg). Diabetic mice were randomized to receive placebo, oral Y1 antagonist BIBO3304 or metformin for 6 weeks. Metabolic and glucose homeostasis parameters were examined thereafter. (B) Non-fasted blood glucose levels at the indicated time points were measured from placebo and BIBO3304 treated mice. n = 8 per group. (C) Six-hour and overnight fasted blood glucose levels. n = 7-8 per group. (D) Urine glucose levels. n = 6-7 per group. (E) Non-fasting blood glucose levels at the indicated time points were measured from placebo, BIBO3304 or metformin treated mice. Results expressed as area under the curve. n = 4-6 per group. (F) Intraperitoneal glucose tolerance tests (1 g/kg body weight) on 6 h fasted diabetic mice treated with placebo or BIBO3304 for 4 weeks. Blood glucose levels during glucose tolerance tests were monitored and results are expressed over the time course and as area under the curve. n = 8 per group. (G-H) Diabetic mice treated with placebo or BIBO3304 were fasted overnight or 6 h and i.p. pyruvate tolerance tests (1 g/kg body weight) or insulin sensitivity tests (0.75 i.u./kg body weight) were performed, respectively. Blood glucose levels during tolerance tests were monitored and results are expressed over the time course and as area under the curve. n = 6-8 per group. (I) Plasma insulin levels throughout intravenous glucose tolerance tests (1 g/kg body weight) from mice treated with placebo or BIBO3304. n = 5-6 per group. (J) C57BL/6 mice were rendered diabetic by multiple high-dose of STZ injections (6 doses, 50 mg/kg body weight). Non-fasted blood glucose levels at the indicated time points were measured from placebo and BIBO3304 treated mice. n = 5-6 per group. (K) Sections of pancreas from placebo or BIBO3304 treated mice were stained for insulin (green) and nuclear counterstained with DAPI (blue). (L-N) Islet number, islet size and islet proportion were determined across three non-consecutive pancreatic sections per mouse and normalized to total pancreas section area. n = 5-6 per group. Data are mean ± SEM. \**P* < 0.05, \*\**P* < 0.01; calculated by unpaired Student’s *t*-test or two-way ANOVA analysis.

Consistent with the improved glycaemic control, glucose tolerance was also significantly improved in mice that received BIBO3304 treatment, which was due to the enhanced *in vivo* insulin secretory response (Figure 3F and 3I). On the other hand, BIBO3304 treatment had no influence on hepatic glucose production or insulin responsiveness *in vivo* or *ex vivo* (Figure 3G, 3H **and** S3I). The improvement in glycaemic control by BIBO3304 treatment was not evident in mice receiving high doses of STZ, a model in which the majority of β-cells are lost (Figure 3J). Importantly, this suggests that the anti-diabetic effect of Y1 receptor antagonist BIBO3304 is dependent on the improvement of the functional β-cell mass. In line with this, there was a significant increase in islet numbers in pancreas from HFD/STZ mice receiving 4-week of BIBO3304 treatment (Figure 3K and 3L), while pancreas weights (Figure S3J) remained comparable. While islets from BIBO3304-treated mice were similar in size to the one in the placebo group, the total islet area was significantly greater in the BIBO3304-treated mice, due to an increase in islet number (Figure 3M **&** N). Collectively, these results suggest Y1 receptor antagonism may be clinically beneficial to improve glycaemic control in T2D by improving functional β-cell mass.

### Y1 receptor antagonism improves insulin responsiveness and β-cell function at various stages of diabetes progression

Genetically diabetic leptin receptor-deficient *db/db* mice are obese, insulin resistant, and display hyperglycaemia at an early age and transition from β-cell compensation to failure with a pathophysiological sequence of events similar to human T2D [19]. To test the effects of BIBO3304 on glycaemic control also in *db/db* mice, we chose an early (4-10 weeks old) and a late (10-16 weeks old) stage of T2D. Four-week-old *db/db* mice were treated daily with BIBO3304 or placebo over a period of 6 weeks. Interestingly, after 6 weeks of treatment, the BIBO3304-treated group showed a significantly lower body weight compared to placebo (Figure 4A). While lean mass remained comparable, the observed reduction in body weight in the BIBO3304-treated group was mostly due to a decrease in fat mass (Figure 4B and 4C). Consistently, the absolute weights of individual fat pads revealed that inguinal fat mass was significantly lower in the BIBO3304-treated group compared to placebo (Figure 4D). The reduction in body weight observed in the BIBO3304-treated group was not due to changes in appetite as evidenced by the absence of a significant difference in food intake (Figure S4A). Importantly, fed and fasted blood glucose were significantly lower in BIBO3304-treated mice compared to placebo (Figure 4E). BIBO3304-treated mice also exhibited significantly lower fasted plasma insulin levels (Figure 4F), suggesting that the inhibition of Y1 receptor signalling may improve glucose control via increasing insulin action. Consistently, insulin tolerance tests revealed that BIBO3304-treated *db/db* mice exhibited a markedly improved insulin responsiveness, as evidenced by lower blood glucose across the duration of 120-minutes and when quantified as area under the curve (Figure 4G). Although the differences were not statistically significant, BIBO3304-treated mice displayed modestly improved whole-body glucose tolerance (Figure S4B and S4C). The enhanced insulin responsiveness was correlated with increased insulin induced Akt phosphorylation in muscle (Figure 4H) but not in the liver or adipose tissue (Figure S4D). In line with this, insulin-induced 2DG glucose uptake was significantly enhanced in extensor digitorum longus (EDL) muscle isolated from *db/db* mice treated with BIBO3304 for 4 weeks, an effect that was impaired in the placebo group (Figure 4I). Strikingly, human muscle *NPY1R* expression was 3-fold higher in obese compared to lean subjects (Figure 4J). The increased *NPY1R* expression in human vastus lateralis muscle also exhibited a positive correlation with BMI (Spearman’s r=-0.6291, p=0.005) as well as fasting blood glucose levels (Spearman’s r=-0.5273, p=0.0245) (Figure 4K and 4L). Consistently, we show in primary human myotubes that insulin-stimulated glucose uptake was suppressed significantly by NPY, an effect that was diminished in the presence of BIBO3304 (Figure 4M). Taken together, these results suggest that on the early onset of T2D, Y1 receptor antagonism attenuates hyperglycaemia which can be attributed to improved insulin action as a consequence of reduced adiposity and/or directly due to inhibition of Y1 receptor in muscle.

**Figure 4.**
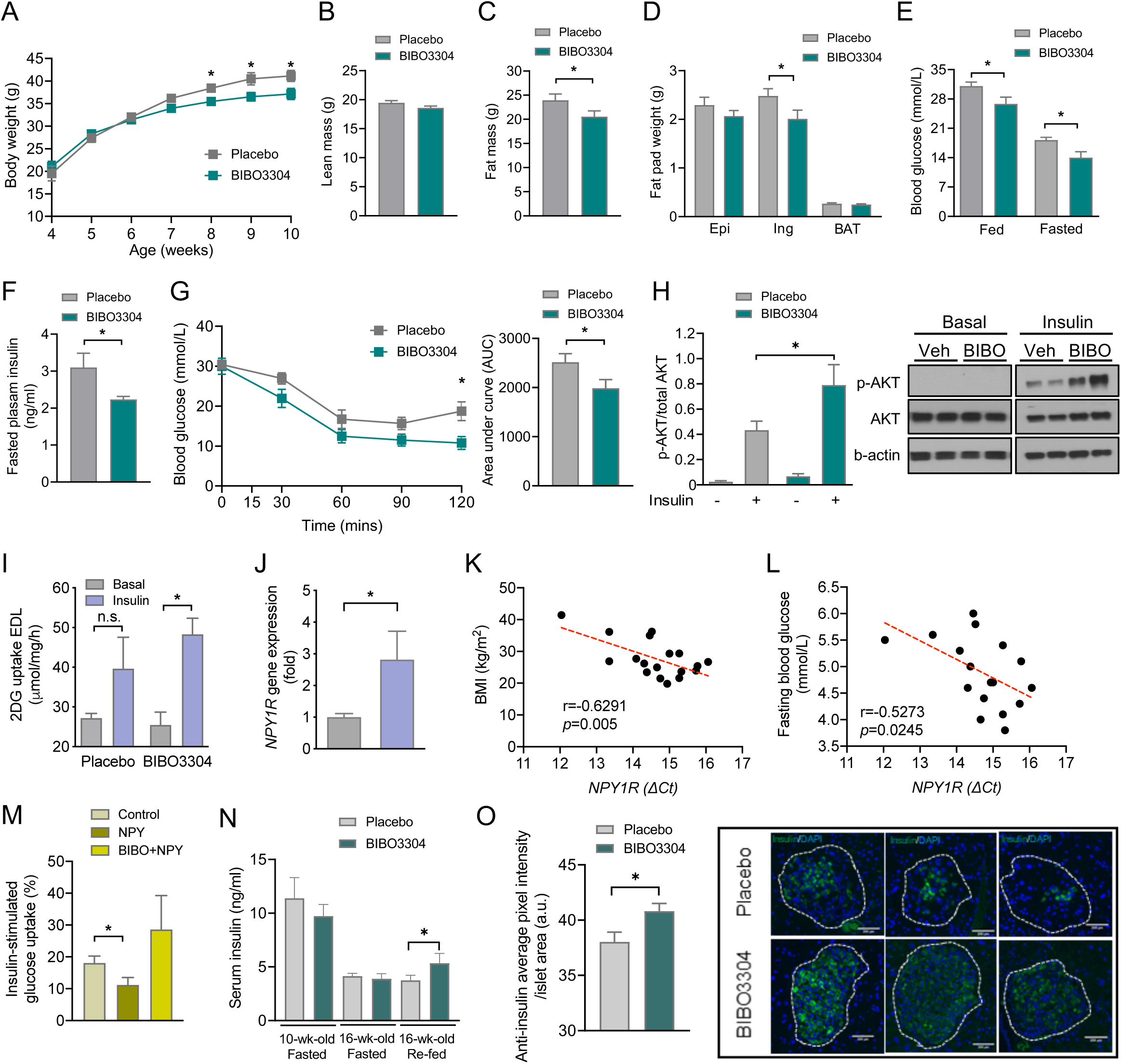
Y1 receptor antagonist BIBO3304 improves hyperglycemia, insulin sensitivity and preserves functional β-cell mass in *db/db* mice. Four-week-old leptin receptor deficient *db/db* mice were randomized to receive placebo or oral Y1 antagonist BIBO3304 for 6 weeks. (A) Weekly body weight of *db/db* mice treated with placebo or oral BIBO3304 (n = 4-6 per group). (B-C) Whole body lean and fat mass as determined by EchoMRI analysis in *db/db* mice treated with placebo or oral BIBO3304 (n = 7 per group). (D) Dissected weights of individual white adipose tissue from epididymal (Epi), inguinal (Ing) and brown adipose tissue (BAT) (n = 4-5 per group). (E) Fed and fasted blood glucose levels in *db/db* mice treated with placebo or oral BIBO3304 (n = 5-6 per group). (F) Fasting plasma insulin levels in *db/db* mice treated with placebo or oral BIBO3304 (n = 6-8 per group). (G) *db/db* mice treated with placebo or BIBO3304 were fasted 6h or overnight and intraperitoneal insulin tolerance tests (2.5 i.u./kg body weight) were performed. Blood glucose levels during tolerance tests were monitored and results are expressed over the time course and as area under the curve. (n = 5 per group). (H-I) EDL muscle isolated from *db/db* mice treated with placebo or BIBO3304, and insulin-stimulated glucose uptake and Akt activation were determined. The muscle homogenates were subjected to SDS-PAGE and western blot analysis using anti-phospho Ser 473 Akt, total Akt and β-actin antibodies (*n* = 5-6). Results shown are a representative blot and quantitative densitometry analysis. The cropped gel is used in the figure and full-length gel is presented in Supplemental Figure S4C. (J) *NPY1R* expression in human muscle from lean (BMI < 25) and overweight/obese (BMI> 25) subjects. Subject numbers: lean = 7 and overweight/obese = 11. (K-L) Correlation between the fasting blood glucose or BMI and the expression of *NPY1R* mRNA (delta CT values) in human muscle of obese and lean control subjects. Total subjects = 18. (M) Primary human muscle cells (n =3) were cultured and insulin-stimulated glucose uptake was determined following the treatment with NPY (Leu31, Pro34) or NPY+Y1 receptor antagonist BIBO3304. Results were presented as percentage increase from basal, and data are the average of 3 independent experiments. (N) Four- and ten-week-old leptin receptor deficient *db/db* mice were randomized to receive placebo or oral Y1 antagonist BIBO3304 for 6 weeks. Fasted and re-fed serum insulin levels were measured (n = 5-8 per group). (O) Pancreases from placebo or BIBO3304 treated mice at 16 weeks of age were weighed and fixed in formalin and processed for immunostaining of insulin (green) and nuclear counterstained with DAPI (blue). Insulin intensity was determined by screening 138 and 172 islets on placebo and BIBO3304 treated pancreatic sections, respectively. Insulin intensity was presented as insulin positive pixel normalized to the islet area. Data are means ± SEM \**P* < 0.05, \*\**P* < 0.01; calculated by unpaired Student’s *t*-test or two-way ANOVA analysis.

While BIBO3304 treatment of late stage diabetic *db/db* mice at 16 weeks of age did not show any effects on body weights, lean mass, fat mass, fat pads mass and insulin response (Figure S4E-S4I), and the impaired β-cell compensation in the 16-week-old *db/db* mice became evident as indicated by a greater than 2.5-fold reduction in serum insulin level as compared to10-week-old *db/db* mice (11.4±1.91 ng/ml in 10-week-old vs. 4.1±0.26 ng/ml in 16-week-old *db/db* mice, n=5-6) (Figure 4N). It is of interest that hyperinsulinemia in the early pathogenesis of T2D was associated with a greater than 60% reduction in *Npy* and *Npy1r* expression in islets of 8-week-old *db/db* mice when compared to the non-diabetic *db/+* mice (Figure S4J), which further supports an inhibitory role for NPY/Y1R signaling in the regulation of insulin secretion. More importantly, BIBO3304 treatment led to a significant enhancement of insulin secretion in response to re-feeding after an overnight fast in the 16-week-old *db/db* cohort when compared to the placebo group (Figure 4N), suggesting an increase in postprandial-induced insulin secretion. To further investigate whether Y1 receptor antagonism has also effects on preserving β-cell mass during the transition from β-cell compensation to failure, pancreases from 16-week-old *db/db* mice treated daily for 6 weeks with BIBO3304 or placebo were examined. Pancreatic histological analysis revealed that pancreas weights, pancreatic islet area, islet number, and islet proportion were comparable between BIBO3304 and placebo treated groups (Figure S4K-S4N). However, a significant increase in the intensity of insulin staining in β-cells was observed in pancreases from BIBO3304 treated *db/db* mice compared to placebo treated *db/db* mice (Figure 4O). Collevtively, these results suggest that Y1 antagonism preserves β-cell insulin content and secretory capacity, thereby delaying diabetes progression in *db/db* mice.

## DISCUSSION

In this study, we demonstrated that the increased NPY and Y1 receptor expression in islets are associated with reduced insulin secretion in human type 2 diabetes. In addition, pharmacological inhibition of Y1 receptor signalling under diabetogenic conditions resulted in improved glucose-stimulated insulin secretion and reduced β-cell death *ex vivo*. Y1 receptor antagonism with BIBO3304 improved β-cell function and preserved functional β-cell mass, thereby resulting in better glycaemic control in HFD/STZ-induced diabetic mouse models. Furthermore, treatment of early-diabetic *db/db* mice with BIBO3304 resulted in reduced adiposity accompanied by lower fasted and postprandial blood glucose levels due to enhanced insulin sensitivity and muscle glucose uptake. Importantly, we also showed that administration of BIBO3304 in severely diabetic *db/db* mice delays diabetes progression through preserving functional β-cell mass during the transition from β-cell compensation to failure. These findings extend our previous studies which revealed that inhibition of Y1 receptor signalling improves β-cell function in both rodent and human islets which can be utlised to improve islet transplantation outcomes and a delayed onset of T1D [20].

Compared to the critical role of neuronal NPY and its receptor Y1 in the regulation of appetite and energy metabolism, the role of NPY-Y1 receptor signalling in the regulation of β-cell function and mass, in particular in human type 2 diabetes, is far less clear. In this study, we demonstrate that in addition to NPY, its receptor Y1 expression was also upregulated in islets from subjects with T2D and the augmented islet NPY and Y1 receptor expression is associated with β-cell dysfunction and failure, thus representing a potential driver of diabetes onset. These results are in line with studies that showed impaired glucose-stimulated insulin secretion in β-cell specific NPY overexpressed islets [21]. Indeed, our results further indicate that pharmacological inhibition of Y1 receptor using BIBO3304 in mouse islets resulted in decreased apoptosis as well as improved β-cell function under various diabetogenic stress conditions. These results demonstrate the significance of NPY-Y1 receptor signalling inhibition, which not only enhances insulin secretion but also protects β-cell against apoptosis.

More importantly, results from these preclinical proof-of-concept studies revealed that Y1 receptor antagonism with BIBO3304 can act as an insulin sensitiser when β-cells remain functioning (early pathogenesis of T2D) and prevents β-cell loss at the late stage of T2D. Y1 receptors are G-protein coupled receptors which preferentially associate with G_i/o_ G-protein and therefore act in an inhibitory fashion [22]. Intracellular cAMP levels are reduced in target cells in response to Y1 receptor ligands, whereas cAMP is increased in response to Y1 antagonism [23]. In line with this, a previous study on islets isolated from Y1 receptor knockout mice found up-regulated cAMP levels [20]. The cAMP signalling-dependent mechanisms have been identified to play a critical role in improving insulin secretion and β-cell survival in diabetes [24]. For instance, pharmacological cAMP inducers such as GLP-1 agonist exendin-4, decreases cytokine- and ER stress-induced impaired β-cell function and apoptosis via a cAMP-dependent signalling pathway in both rodent and human β-cells [25, 26]. Supporting this notion, our findings showed that under diabetogenic conditions, islets treated with BIBO3304 exhibited significant improvement in glucose-stimulated insulin secretion, suggesting that this is attributed to enhanced intracellular cAMP levels.

Most of the known effects of the NPY system in the development of obesity arise from the central activation of Y1 receptors, where it plays a critical role in the regulation of appetite and energy homeostasis [9]. Inhibition of Y1 receptors or NPY deficiency in the brain have been linked to decreased body weight gain and adiposity by decreasing energy intake and increasing energy expenditure [27, 28]. Our findings revealed that the administration of the non-brain penetrable Y1 receptor antagonist BIBO3304 also resulted in decreased body weight and fat mass in *db/db* mice in the absence of any alteration in food intake, suggesting that BIBO3304 reduces adiposity by acting on mechanisms other than regulation of appetite centrally. This is consistent with results from a previous study conducted by Zhang et al., where it was revealed that conditional knockdown of Y1 receptors in the periphery exhibited a phenotype of reduced RER, indicating increased lipid oxidation [29]. The underlying mechanisms behind the increased lipid oxidation under peripheral Y1 antagonism was reportedly associated with increased levels of carnitine palmitoyltransferase-1 (CPT-1) and upregulation of key enzymes involved in β-oxidation, consequently increasing the capacity of the mitochondria for lipid oxidation and transport of fatty acids particularly in the liver and muscle [29].

In addition to reduced adiposity, BIBO3304 treatment in *db/db* mice also significantly enhanced insulin responsiveness as demonstrated by increased insulin induced AKT phosphorylation and insulin-stimulated glucose uptake in skeletal muscle of *db/db* mice and in primary human muscle cells. The insulin sensitising effect observed in *db/db* mice might be, at least in part, due to reduced body weight and adiposity or muscle fat content. In addition, in line with our finding in primary human myotubes, previous studies showed that deficiency of peripheral Y1 receptor results in increased mitochondrial capacity in muscle [29], supporting a role of Y1 receptor antagonism acting directly on muscle insulin receptor signalling. Nonetheless, these results are consistent with the notion that increasing muscle glucose uptake improves glycaemic control and suggesting that, in addition to reduced adiposity, these effects may at least in part be responsible for the observed improvement in glucose homeostasis in BIBO3304 treated *db/db* mice.

In summary, one unmet clinical need in treating T2D is the availability of therapeutics that improves glycaemic control by targeting the underlying β-cell dysfunction and failure. As such the reduced adiposity and improved insulin action seen by the inhibition of NPY/Y1 signalling *in vivo* highlights a potential therapy of targeting these peripheral Y1 receptor pathways to preserve functional β-cell mass, which may ultimately provide greater therapeutic benefits in controlling glucose levels in T2D.

## MATERIALS and METHODS

### Key resources table

| REAGENT or RESOURCE | SOURCE | IDENTIFIER |
| --- | --- | --- |
| Antibodies |  |  |
| Mouse monoclonal anti-ATK (pan) (40D4) | Cell Signaling Technology | Cat#: 2920S |
| Rabbit monoclonal anti-phospho-AKT (Ser 473) | Cell Signaling Technology | Cat#: 9271S |
| Rabbit monoclonal anti β-Actin | Cell Signaling Technology | Cat #: 4970; RRID: AB_2223172 |
| Rabbit monoclonal anti-cleaved caspase-3 (Asp175)(5A1) | Cell Signaling Technology | Cat #: 9664S |
| Rabbit monoclonal anti-BIM (C34C5) | Cell Signaling Technology | Cat #: 2933T |
| goat anti-rabbit-HRP antibody | Agilent Technologies | Cat#: P0448 |
| Goat anti-mouse immunoglobulins/HRP | Agilent Technologies | Cat#:P0447 |
| Guinea Pig Polyclonal Anti-Insulin | Agilent Technologies | Cat #: IR00261-2 |
| Alexa Fluor 488 Goat Anti-Guine Pig IgG (H+L) | Life Technologies | A-11073 |
| Alexa Fluor 594 Goat anti-Rabbit IgG (H+L) | Life Technologies | A11012 |
| Chemicals, Peptides, and Recombinant Proteins |  |  |
| RPMI-1640 cell culture media | Sigma-Aldrich |  |
| streptozotocin | Sigma-Aldrich | Cat #: S0130 |
| 1,1-Dimethylbiguanide hydrochloride (Metformin) | Sigma-Aldrich | Cat #: D150959 |
| BIBO3304 trifluoroacetate | Tocris | Cat #: 2412 |
| Actrapid (human, Insulin) | Novo Nordisk Pharmaceuticals | Cat #:169625 |
| Critical Commercial Assays |  |  |
| Mouse Insulin ELISA | ALPCO Diagnostics | Cat#: AP80-INSMS-E10 |
| Glucose oxidase assay | ThermoFisher | Cat#: A22189 |
| Experimental Models: Organisms/Strains |  |  |
| C57BL/6J |  | C57BL/6J |
| B6.BKS(D)Lepr<sup>db</sup>/J (*db/db*) |  | *db/db* |
| Oligonucleotides |  |  |
| *NPY* TaqMan® Gene Expression Assay | ThermoFisher | Cat #: Hs00173470_m1 |
| *PYY* TaqMan® Gene Expression Assay | ThermoFisher | Cat #: Hs00373890_g1 |
| *PPY* TaqMan® Gene Expression Assay | ThermoFisher | Cat #: Hs00358111_g1 |
| *NPY1R* TaqMan® Gene Expression Assay | ThermoFisher | Cat #: Hs00702150_s1 |
| *NPY2R* TaqMan® Gene Expression Assay | ThermoFisher | Cat #: Hs01921296_s1 |
| *PPYR1* TaqMan® Gene Expression Assay | ThermoFisher | Cat #: Hs00275980_s1 |
| *NPY5R* TaqMan® Gene Expression Assay | ThermoFisher | Cat #: Hs01883189_s1 |
| *RPLP0* TaqMan® Gene Expression Assay | ThermoFisher | Cat #: Hs99999902_m1 |
| *GAPDH* TaqMan® Gene Expression Assay | ThermoFisher | Cat #: Hs99999905_m1 |
| *Ppia* TaqMan® Gene Expression Assay | ThermoFisher | Cat #: Mm02342430_g1 |
| *Bcl2* TaqMan® Gene Expression Assay | ThermoFisher | Cat #: Mm00477631_m1 |
| *Bak1* TaqMan® Gene Expression Assay | ThermoFisher | Cat #: Mm00432045_m1 |
| AHLIKIN, Lepr TaqMan SNP Assay | ThermoFisher | Cat #: 4332077 |
| Software and Algorithms |  |  |
| Prism 8.0 | Graphpad | https://www.graphpad.com/scientific-software/prism/ |
| Image J | GNU General Public License |  |

### Resource availability

#### Lead contact

Further information and requests for reagents may be directed to lead author Dr Chieh-Hsin Yang

#### Materials availability

This study did not generate new unique reagents.

#### Data and code availability

This study did not generate any unique datasets or code.

### Experimental models and subject details

We obtained approval for performing human islet studies from St Vincent’s Institute of Medical Research and St. Vincent’s Clinical School Human Research Ethics Committee. Consent for use of the islets for research was given by relatives of the donors. All mice care and experiments were performed in accordance with protocols approved by the Animal Ethics Committee at St Vincent’s Hospital (AEC No. GBNML 760 and 016/19). Eight-week-old mice were fed a standard chow diet (6% fat) or a high fat diet (23% fat; 45% of total energy from fat; SF04-027; Specialty Feeds) as indicated. B6.BKS(D)Lepr ^db^/J (*db/+*) heterozygous mice were kindly provided by A/Prof Ross Laybutt from Garvan Institute of Medical Research (Sydney, NSW, Australia). Routine genotyping for homozygous *db/db* mice was conducted using TaqMan SNP genotyping assay (AHLIKIN, ThermoFisher Scientific). The eight-week-old C57BL/6 male mice were purchased from The Walter and Eliza Hall Institute (Victoria, Australia) for all in vivo studies. To induce T2D, C57BL/6 mice were fed a high fat diet (SF04-027; Specialty Feeds) for 4 weeks from 8 weeks of age, followed by multiple intraperitoneal injections of low-dose streptozotocin (35 mg/kg) (Sigma Aldrich). Streptozotocin was prepared fresh each time in 0.1 M sodium citrate buffer (pH 4.5) and filter sterilised prior to use. Blood glucose level was measured twice a week until blood glucose reached 15 mmol/L and above. C57BL/6 mice used in islets experiments were bred in house at BioResources Centre (St Vincent’s Hospital). All mice were housed in a temperature-controlled room of 22 ֯C on a 12 h/12 h light/dark cycle (lights on from 0700-1900 hours) with free access to water and food.

### Method details

#### Treatment with Y1 receptor antagonist BIBO3304 and metformin

A non-brain penetrable Y1 receptor antagonist BIBO3304 (Tocris Bioscience) was prepared in Milli Q water at a concentration of 1 mg/ml. C57BL/6 mice (average weight 27.4 ± 0.3 g) were received 0.5 mg BIBO3304 daily in jelly containing 4.9% (wt/v) gelatine and 7.5% (v/v) chocolate flavouring essence as described previously [13]. The obese *db/db* mice at 4- (average weight 20.3 ± 0.9 g) or 12-weeks of age (average weight 42 ± 1.3 g) were given 2.5 mg BIBO3304 once daily via oral gavage, while control mice on placebo treatment received the same volume of Milli Q water. Metformin (Sigma Aldrich) was prepared in Milli Q water, and 0.25 g/kg was given daily via oral gavage. The duration of treatment is as stated in the text for each procedure.

#### Metabolic assessment and body composition measures

The effect of Y1 receptor antagonist BIBO3304 on blood glucose control and body weight were monitored weekly on the same day of the week between 09:00 hours and 10:00 hours. Random blood glucose was taken from tail tipping and measured on an Accu-Check Performa glucometer (Roche, Switzerland). For the fast-refeeding experiment, food was removed from the mice at the dark cycle before the experiment. Blood was collected by retro-orbital bleed after a 16 h fast as well as 30 minutes after refeeding to determine blood glucose and plasma insulin levels. Food intake was measured at the same time points (n = 8 per group). Whole body lean mass and fat mass were measured at the end of study using the whole-body composition analyzer, EchoMRI (Houston, USA).

#### *In vivo* assessment of glucose, insulin and pyruvate tolerance tests

Glucose tolerance tests were performed on 6 h-fasted HFD/STZ mice or overnight fasted *db/db* mice by intraperitoneal injection of 1 g/kg and 0.5 g/kg glucose, respectively. Insulin tolerance was measured by intraperitoneal injection of 0.75 i.u./kg and 2.5 i.u./kg human insulin (Actrapid, Novo Nordisk Pharmaceuticals) on HFD/STZ mice and *db/db* mice after a 6h-fast. Pyruvate tolerance tests were conducted on mice after an overnight fast with intraperitoneal injection of 1 g/kg sodium pyruvate. Blood glucose was measured at basal and, 15, 30, 60, 90 and 120 minutes following glucose, insulin or pyruvate administration. The *in vivo* glucose-stimulated insulin secretion was determined by intravenous glucose tolerance test using 1 g/kg glucose on overnight fasted HFD/STZ mice as previously described. Briefly, mice were anaesthetised and jugular venous catheters were inserted. Mice were allowed to recover for 20 minutes after surgery. A bolus of glucose was given via catheter and blood glucose was measured at 2, 5, 10, 15 and 30 minutes post glucose administration.

#### Pancreatic islet isolation and culture *ex vivo*

Mouse islets were isolated from C57BL/6 and *db/db* mice as previously described [30]. Briefly, Collagenase P (0.45 mg/mL) (Sigma Aldrich) was injected into the bile duct to distend the pancreas. After perfusion, pancreas was excised and incubated at 37°C for 15 minutes. Islets was further purified using Histopaque-1077 gradient (Sigma Aldrich). The isolated mouse islets were cultured in Connaught Medical Research Laboratories (CMRL) 1066 medium (Invitrogen, Life Technologies, Carlsbad, CA, USA) supplemented with 10% fatal calf serum, 100 U/ml penicillin, 100 mg/ml streptomycin and 2 mmol/L l-glutamine. Isolated islets were incubated in 37°C, 5% CO_2_ humidified incubator.

#### Human islet isolation and culture *ex vivo*

Pancreases were obtained from heart-beating, brain-dead donors with consent from next-of-kin and research approval from the St Vincent’s Hospital, Melbourne (HREC-011-04). Human islets were purified using Ficoll density gradients [31] and cultured in Connaught Medical Research Laboratories (CMRL) 1066 medium (Invitrogen, Life Technologies, Carlsbad, CA, USA) supplemented with 10% fetal calf serum, 100 U/ml penicillin, 100 mg/ml streptomycin and 2 mmol/l l-glutamine. All islets were incubated in a 37°C, 5% CO_2_ humidified incubator. Insulin stimulation index was determined and presented as the ratio of insulin secretion at 28 mmol/l to that of at 2.8 mmol/l from the same islets.

#### Skeletal muscle biopsies from human donors

Eighteen non-diabetic males with an average of 40.4 ± 3.9 years of age, Body Mass Index (BMI) 27.7 ± 1.4 kg/m^2^, and fasting blood glucose of 4.9 ± 0.15 mmol/L were included. Muscle samples were acquired from the *vastus lateralis* under local anesthesia (Xylocaine 1%) using the percutaneous needle biopsy technique with suction. The samples were snap frozen in liquid nitrogen and stored at −80°C until analyses. The methods for participant recruitment and muscle biopsy were approved by the Human Research Ethics Committee, Victoria University.

#### Glucose-stimulated insulin secretion in isolated islets

Wild-type C57BL/6 islets were incubated in the respective diabetogenic stressors for the indicative duration: Inflammation: islets were incubated with proinflammatory cytokine cocktail (50 ng/ml TNFa, 250 ng/ml IFNg and 25 ng/ml IL1b) for 48 hours; oxidative stress: 10 mM H_2_O_2_ for 16 hours; ER stress: 1 mM thapsigargin for 24 hours. Following culture, islets were handpicked and pre-incubated for 1 hour in HEPES-buffered-KREBS buffer containing 0.2% BSA and 2.8 mmol/L D-glucose. Subsequently, 15 size matched islets were incubated at 37°C for another 1 hour in KREBS buffer containing either 2.8 mmol/L or 20 mmol/L D-glucose, treated with or without 1 μM BIBO3304. Culture medium was collected, and insulin secretion was assayed using a mouse insulin ELISA kit (ALPCO Diagnostics, Salem, NH, USA).

#### DNA fragmentation assay

To induced islet cell death, wild-type C57BL/6 islets were incubated in the respective diabetogenic stressors: Inflammation: islets were incubated with proinflammatory cytokine cocktail (50 ng/ml TNFa, 250 ng/ml IFNg and 25 ng/ml IL1b) for 72 hours; oxidative stress: 70 mM H_2_O_2_ for 18 hours; ER stress: 5 mM thapsigargin for 72 hours, glucolipotoxicity: 25 mmol/L glucose plus 0.5 mM palmitate for 96 hours. Cell apoptosis was measured by analysis of DNA fragmentation as described previously [32]. Briefly, islets in uniform size were handpicked into 3.5 cm Petri dishes containing the appropriate stimuli to induce apoptosis in 1.5 ml complete CMRL medium. At the end of the culture period, islets were dispersed by trypsin digestion for 5 min at 37°C, followed by mechanical disruption by pipetting up and down for 10 times. The dispersed islet cells were then resuspended in 150 ml of Nicoletti buffer containing 50 mg/ml propidium iodide (Miltenyi Biotec), 0.1% (wt/v) sodium citrate and 0.1% (v/v) Triton X-100 [33]. The cells were then analyzed on a LSRFortessa Flow Cytometer (Becton Dickinson, Franklin Lakes, NJ). Cells undergoing apoptosis were identified by their apparent sub-diploid DNA content as reported previously [34].

#### 2-Deoxyglucose uptake measurement in EDL muscle

Db/db mice were fasted overnight then euthanised using CO_2_ chamber. Left and right EDL muscles were quickly excised and bathed in carbogenated Krebs-Henseleit buffer (KHB) (119 mM NaCl, 4.7 mM KCl, 2.5 mM CaCl_2_, 1.2 mM MgSO_4_, 1.2 mM KH_2_PO_4_, 25 mM NaHCO_3_, pH 7.4, 30 ⁰C) with constant shaking. After 30 min of pre-incubation, muscles were transferred to fresh carbogenated KHB containing 10 mU/mL insulin (or KHB without insulin as control) for 30 min. Subsequently, muscles were transferred to fresh KHB containing 2 mM 2-Deoxy-d-[1,2-3H]-glucose (0.15 μCi/mL) and 16 mM d-[1-^14^C] mannitol (0.1 μCi/mL) for 15 min. After the incubation, muscles were rapidly rinsed with ice cold KHB buffer, then snap frozen in liquid nitrogen and stored at −80 ⁰C. Muscle samples were next lysed in ice-cold radioimmunoprecipitation assay (RIPA) buffer (400 μL/muscle) with protease and phosphatase inhibitor cocktail (Cell Signalling) using TissueLyser II (QIAGEN). Half of the lysate was mixed with scintillation cocktail for scintillation counting using Liquid Scintillation Analyzer (PerkinElmer), and the other half was used for immunoblotting.

#### 2-Deoxyglucose uptake measurement in human myotubes

Three lines of primary human myoblasts originated from skeletal muscle samples of three non-diabetic male participants (age: 64, 72, and 80 years) were used to assess insulin-stimulated glucose uptake. Myogenic differentiation of myoblasts was initiated when cells grew to ∼80% confluence in 12-well plates, the growth medium (10 % fetal bovine serum [FBS] in α-MEM) was replaced with the differentiation medium containing 2 % horse serum in α-MEM. After 5 days of differentiation (Differentiation Medium was refreshed every other day), cells were treated with/without 0.5 μM NPY and/or 1 μM BIBO3304 in serum-free medium for 24 h. Following the treatment, cells were stimulated with 100 nM insulin (or without insulin) in Glucose Uptake Buffer (GUB) (10 mM HEPES, 2.5 mM NaH_2_PO_4_, 150 mM NaCl, 5 mM KCl, 1.2 mM CaCl_2_, 1.2 mM MgSO_4_, 0.1% BSA, pH7.4) for 45 min. In the last 15 min of stimulation, 1 mM 2-Deoxy-d-[1,2-^3^H]-glucose (1 μCi/mL) was spiked into the GUB. After the incubation, cells were washed three times with ice-cold PBS, then lysed with 0.1 M NaOH (200 μl/well). 150 μl of the lysate was pipetted into vials with scintillation cocktail for scintillation counting, and the remaining was used in protein assay for normalisation purpose.

#### RNA extraction and quantitative real-time PCR

Total RNA of mouse islets was extracted using RNeasy Plus Mini Kit (Qiagen). Other tissues including mouse liver, muscle and adipose tissues were excised and snap frozen in liquid nitrogen, and RNA was isolated using RNAzol Reagent (Sigma, St. Louis, MO) following the manufacturer’s instructions. Isolated mRNA was reverse transcribed into cDNA using the Superscript IV First-Strand Synthesis System (Invitrogen, Australia) and performed quantitative real-time PCR with the Light-Cycler 480 Real-Time PCR system (Roche, Switzerland). Relative gene expression of NPY ligands and receptors was performed under the assumption that the probes binding efficiency are equal. Human *RPLP0*, *GAPDH* and mouse *Ppia* were used as housekeeping genes for normalisation as indicated in the text. Primer details are listed in the Supplementary Table 2. The amplification condition used in all the RT-qPCR experiment was: 95 °C for 10 min, 95 °C for 15 s, 60 °C for 60 s for 40 cycles. Relative quantification was determined using the 2^−ΔΔCt^ method.

#### Immunoblotting

200 islets per sample from *db/db* or *db/+* mice were cultured in 3 mL of complete CMRL medium and treated with or without BIBO3304 (1 μM) for 36 hours. Islets were lysed in ice-cold RIPA lysis buffer supplemented with protease and phosphatase inhibitor cocktails (Cell Signalling Technology). Protein concentrations were determined with BCA protein assay (Pierce, Thermo Fisher Scientific). Proteins were resolved in SDS-PAGE gel (4-20% gradient polyacrylamide gel electrophoresis, Mini-PROTEAN® Precast Gels, Biorad). Blots were blocked for 1 hour with 5% non-fat dry milk in PBS/0.1% tween-20 (Sigma Aldrich), and subsequently incubated overnight at 4 °C with respective primary antibodies: pan-AKT antibody (40D4) (1:2,000; 2920S; Cell Signaling Technology), phospho-AKT (Ser473) (1:1,000; 9271S; Cell Signaling Technology), BIM (C34C5)(1:1,000; 2933T; Cell Signaling Technology), cleaved caspas-3 (Asp175)(5A1E) (1:1,000; 9664S; Cell Signaling Technology) or β-actin (1:2,000; 4970; Cell Signaling Technology). Following incubation, membranes were washed 3×10 minutes in PBS-T, then incubated with HRP-linked secondary antibodies for 1 hour in 5% milk in PBS-T at room temperature. After 3×10 minutes washes, immunoreactive signals were visualised using SuperSignal™ West Femto Maximum Sensitivity Substrate (Thermo Fisher Scientific), then developed using Super RX Fuji X-ray film (Fujifilm, Tokyo Japan). Protein band intensities were quantified using Image J. Cleaved caspase-3 and BIM protein signals were normalised against tubulin as a loading control.

#### Immunofluorescent staining on pancreatic histochemical analysis

Whole pancreas was excised and fixed in 4% PBS-buffered paraformaldehyde overnight at room temperature and embedded into paraffin. Slides with 5 μm thick pancreas sections were deparaffinized using Histolene (Trajan Scientific, Australia), rehydrated with ethanol (100%, 100%, 90%, 70%), and blocked in 10% FBS in PBS for 30 minutes at room temperature. Subsequently, sections were incubated for 2 hours at room temperature with polyclonal guinea pig anti-insulin antibody (1:5, Agilent Technologies). Slides were then washed 3 x 5 minutes with PBS and incubated with the Anti-guinea pig IgG Alexa Fluor 488 (1:200, Life Technologies) diluted in 10% FBS for 1 hour at room temperature. The resulting slides were then mounted in a mounting medium containing DAPI. Slides were scanned at 20x magnification using 3D Histech Panoramic SCAN II slide Scanner (Phenomics Australia Histopathology and Slide Scanning Service, University of Melbourne). For β-cell mass measurement, Islets were outlined manually on the digital images. Islet area and islet number were analysed using digital image processing software Image Scope (Aperio). Two sections separated by at least 150 μm was used for each mouse (n=8 per treatment). Cell mass of pancreatic β-cells was determined as the product of wet pancreas weight and the ratio of insulin positive/total pancreas area.

#### Hepatic glucose production assay

Primary hepatocytes were isolated from C57BL/6 mice at 8 weeks of age and plated at a density of 1 x10^6^ cells in 6-well plates with the plating medium (Williams’ E medium supplemented with 10% fetal bovine serum, 1% penicillin-streptomycin and 1% of L-glutamine) for 4 hours followed by starvation overnight in low glucose DMEM supplemented 1% L-glutamine and 1% Penicillin-Streptomycin. The following day, cells were pre-treated treated with/without 0.5 mM NPY and/or 1 mM BIBO3304 for 1 hour. Subsequently, the cells were washed once with PBS, and the assay medium (DMEM without glucose, 1% penicillin-streptomycin, 2 mM of sodium pyruvate, 20 mM sodium lactate, pH 7.4) was added with/without 0.5 mM NPY and/or 1 mM BIBO3304 for 6 hours. Glucose production was assayed with the Amplex Red glucose assay kit (Invitrogen), and cell lysate was used in protein assay for normalisation.

#### Statistical analysis

All data are presented as mean ± SEM. A Student’s *t*-test was conducted to test difference between two groups of mice. Restricted randomisation was used to achieve treatment group with similar numbers of mice. Sample size was estimated on previously published studies of our and other’s research groups [13, 14, 35, 36]. Differences among groups of mice were assessed by two-way ANOVA or repeated-measures ANOVA. Correlation coefficient was calculated using Spearman’s rank correlation coefficient. Statistical analyses were assessed using Prism software 8.0. All experiments requiring the use of animals or animals to derive cells were subject to randomization based on litter. Differences were regarded as statistically significant if \**P* < 0.05; \*\**P* < 0.01; \*\*\**P* < 0.001; \*\*\*\**P* < 0.0001.

## ACKNOWLEDGEMENTS

This work was supported by the National Health and Medical Research Council (NHMRC) of Australia in the form of a project grant #1158242 to KL. This work was also supported by a Diabetes Australia Project grant (Y19G-LOHK) and Australia Diabetes Society Skip-Martin Fellowships to KL. Supported in part by the Victorian Government’s Operational Infrastructure Support Program. We thank all organ donors and their families, Donatelife and the staff of St Vincent’s Institute involved in the human islet isolation program.

## AUTHOR CONTRIBUTIONS

C.H.Y, D.A.O, X.Z.L, S.N, S.F and E.P, designed and performed research and contributed discussion, C.H.Y, J.W.S, J.O, S.G, Y.S contributed discussion and reviewed manuscript. A.M-A and C.S contributed to research experiments and reviewed manuscript. T.L, I.L, R.D.L, H.H contributed discussion and edited manuscript. H.E.T and K.L contributed discussion, wrote manuscript, reviewed/edited manuscript. All authors read and approved the final manuscript.

## THE PAPER EXPLAINED

### Problem

Loss of functional β-cell mass is a key factor contributing to poor glycaemic control in advanced type 2 diabetes. Hence, one of the most pressing unmet medical needs in type 2 diabetes is the development of new therapeutics that provide β-cell protective effects, such as improvement of β-cell function, mass and survival. Improved understanding of diabetes pathophysiology and the identification of a new biochemical pathway that regulates β-cell function and mass will be extremely valuable for the development of more effective therapeutic approaches for diabetes.

### Results

In this study, we demonstrated that the increased NPY and Y1 receptor expression in islets from patient with type 2 diabetes correlated with reduced β-cell function. Importantly, in a preclinical study, pharmacological inhibition of neuropeptide Y1 receptors by BIBO3304, a selective orally bioavailable neuropeptide Y1 receptor antagonist, significantly improved β-cell function and preserved β-cell mass, thereby resulting in better glycaemic control. Furthermore, Y1 receptor antagonist BIBO3304 exhibited similar efficacy to attenuate hyperglycaemia when compared with a first-line oral anti-diabetic drug, metformin. Collectively, these results demonstrate that inhibition of Y1 receptor by BIBO3304 represents a potential β-cell protective therapy for improving functional β-cell mass and glycaemic control in type 2 diabetes.

### Impacts

This research is the first to uncover a novel causal link of increased islet NPY-Y1 receptor signaling to β-cell dysfunction and failure in human type 2 diabetes, contributing to the understanding of the pathophysiology of type 2 diabetes. These novel findings provide preclinical proof-of-concept for improving functional β-cell mass and resulting in better glycaemic control by targeting the NPY-Y1 receptor pathway. Findings from the current studies provide a significant conceptual advance that could have translational potential for improving treatment of type 2 diabetes.

## SUPPLEMENTAL FIGURE LEGENDS

**Supplemental Figure 1.**
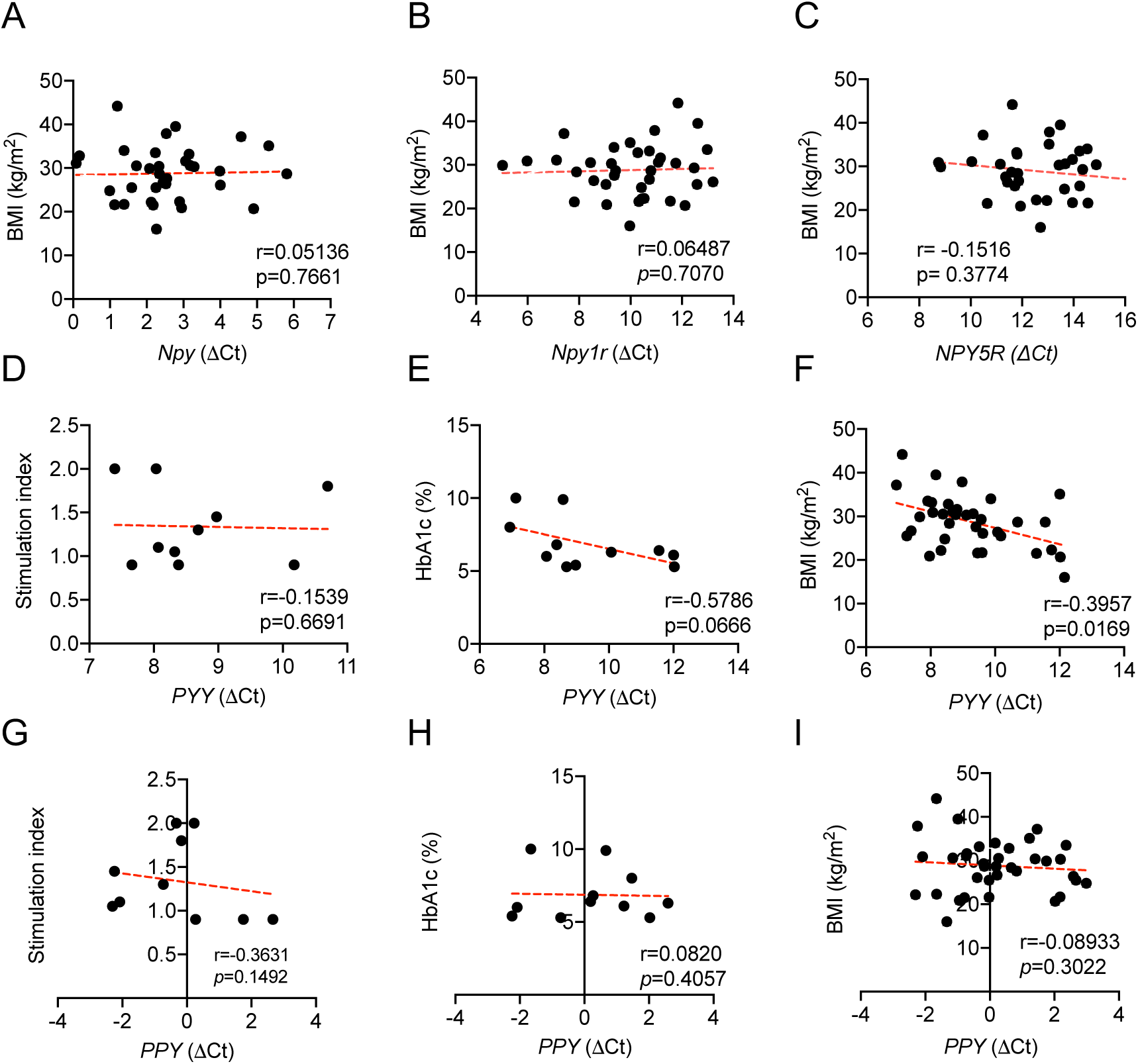
Correlation between the stimulation index, HbA1c or BMI and the islet NPY system expression in type 2 diabetes. (A-C) Correlation between BMI and the expression of *NPY*, *NPY1R* and *NPY5R* mRNA in human islets of subjects with type 2 diabetes and non-diabetic control subjects. Total subjects = 34. (D-F) Correlation between the insulin stimulation index, BMI or HbA1C and the expression of *PYY* mRNA in human islets of subjects with type 2 diabetes and non-diabetic control subjects. Analysis was done with a total number of 9 subjects for stimulation index, 36 subjects for BMI and 11 subjects for HbA1C. (G-I) Correlation between the insulin stimulation index, BMI or HbA1C and the expression of *PPY* mRNA in human islets of subjects with type 2 diabetes and non-diabetic control subjects. Total number of 10 subjects in stimulation index, 36 subjects in BMI and 11 subjects in HbA1C. Data are mean ± SEM. *P* values by two-tailed Spearman’s correlation analysis.

**Supplemental Figure 2.**
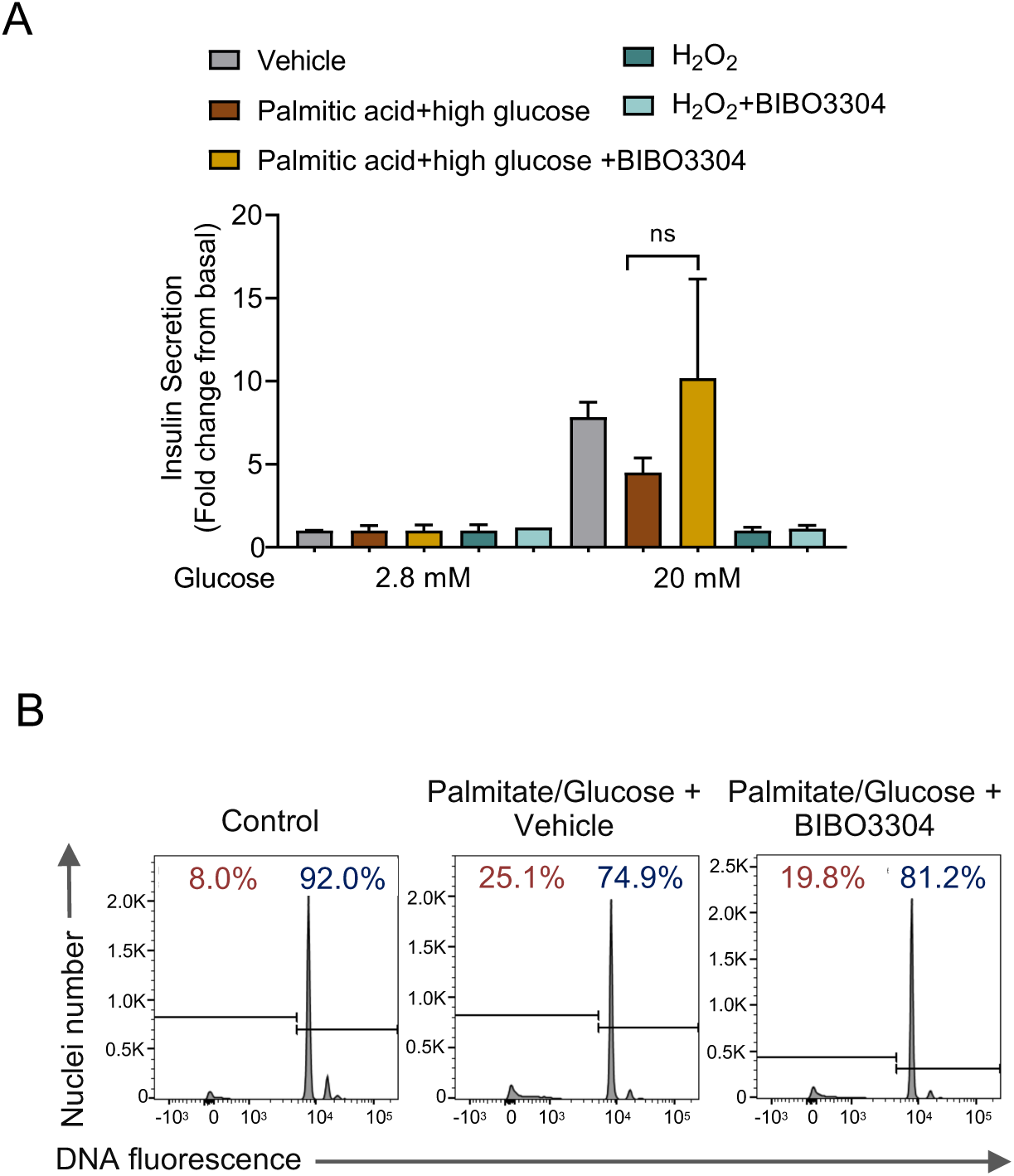
Glucose stimulated insulin secretion and cell death analysis in Y1 receptor antagonist treated islets under glucolipotoxicity and oxidative stress conditions. (A) Pancreatic islets from C57BL/6 mice were isolated and cultured. Islets were exposed to 25 mmol/L glucose and 0.5 mM palmitate ± 1 μM of BIBO3304 for 96h or 10 μM H_2_O_2_ ± 1 μM of BIBO3304 for 16h (*n* = 3). Glucose-stimulated insulin secretion was determined in response to 2.8 and 20 mmol/L glucose. (B) DNA fragmentation in response to glucolipotoxicity (25 mmol/L glucose and 0.5 mM palmitate ± 1 μM of BIBO3304 for 72h) was measured by flow cytometry. Representative FACS profiles are shown and the results are representative of islets from a minimum of n=3 individual mice per group. Data are mean ± SEM. \**P* < 0.05, \*\**P* < 0.01, calculated by unpaired Student’s *t*-test.

**Supplemental Figure 3.**
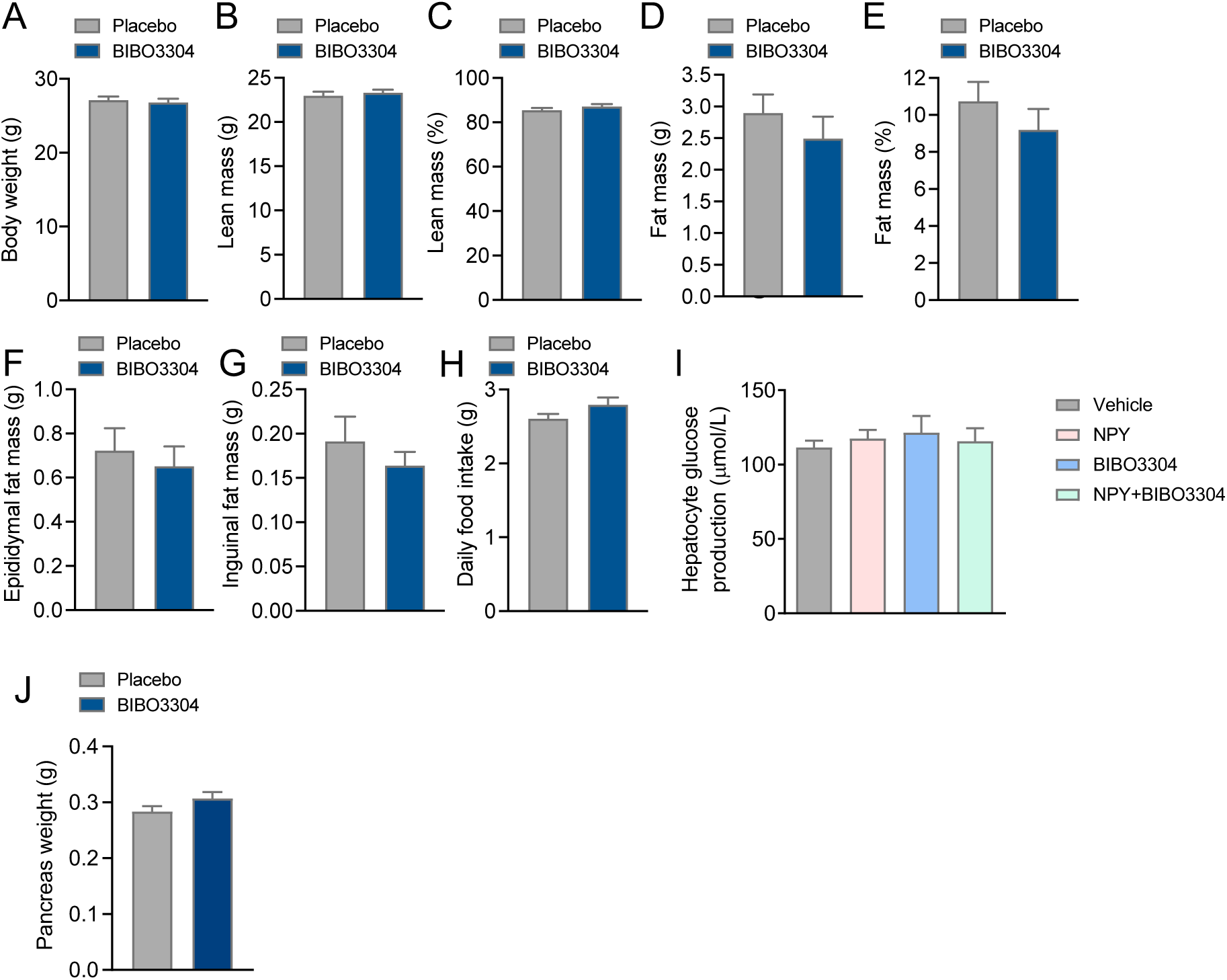
Y1 receptor antagonist BIBO3304 treatment did not alter adiposity, food intake or hepatic glucose production in HFD/STZ-induced diabetes mice. C57BL/6 mice were fed a high fat diet for 4 weeks and rendered diabetic by multiple low-dose STZ injections (6 doses, 35mg/kg). Diabetic mice were randomized to receive placebo or oral Y1 antagonist BIBO3304 for 4 weeks. Metabolic and glucose homeostasis parameters were examined thereafter. (A) Body weight of HFD/STZ-induced diabetes mice treated with placebo or oral BIBO3304 (n = 6-8 per group). (B-E) Whole body lean and fat mass as determined by EchoMRI analysis in placebo or oral BIBO3304 treated HF/STZ mice (n = 6-8 per group). (F-Dissected weights of individual white adipose tissue from epididymal (Epi) and inguinal (Ing) (n = 6-8 per group). (H) Daily food intake of HFD/STZ-induced diabetes mice treated with placebo or oral BIBO3304 (n = 6-8 per group). (I) Hepatocytes were isolated and glucose production was performed in the presence of NPY, BIBO3304 or NPY+BIBO3304 (n = 6-8 per group). (J) Dissected weights of pancreas from HFD/STZ-induced diabetes mice treated with placebo or oral BIBO3304 (n = 6-8 per group). Data are means ± SEM. \**P* < 0.05, \*\**P* < 0.01; calculated by unpaired Student’s *t*-test.

**Supplemental Figure 4.**
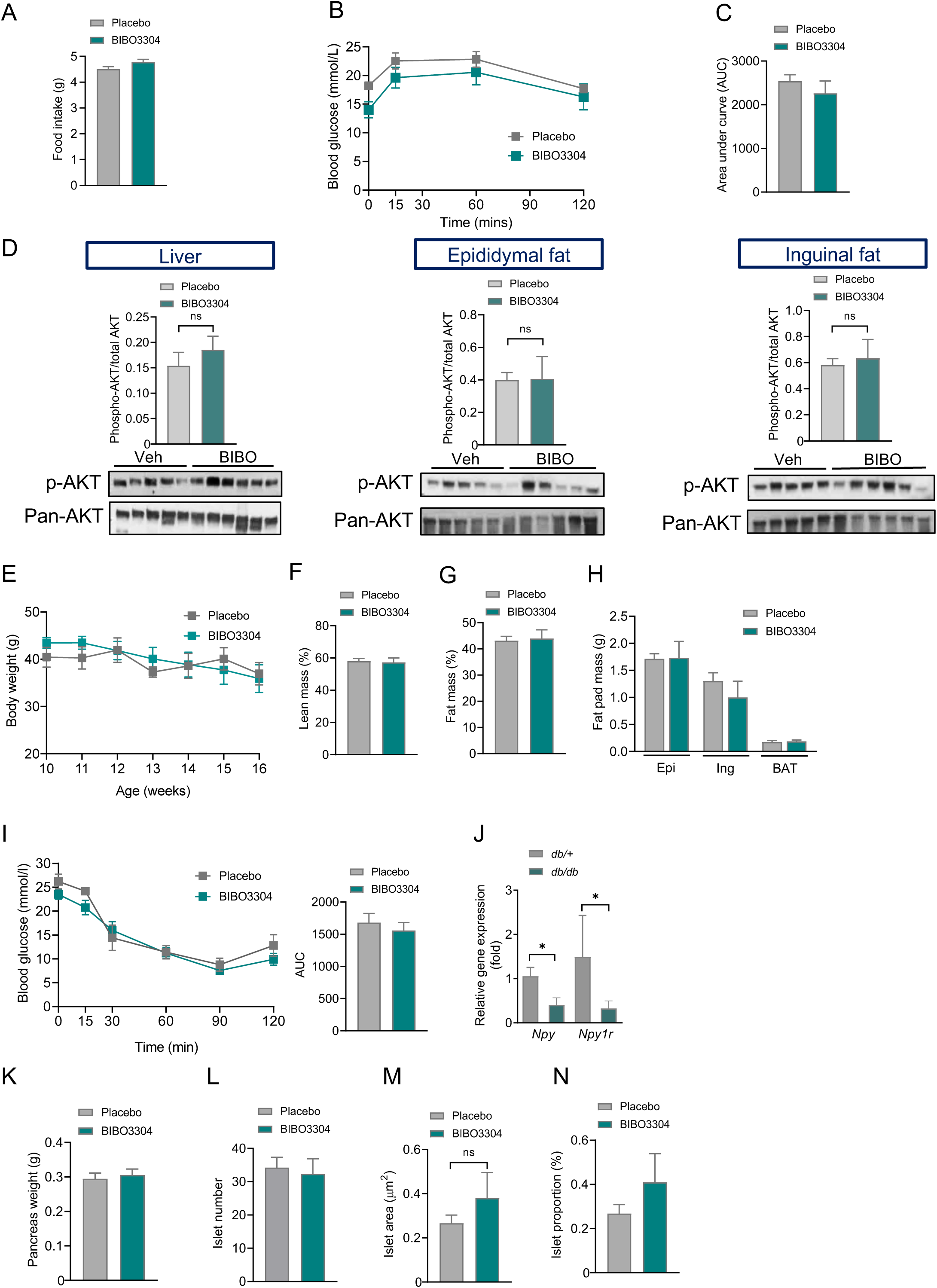
Metabolic and glucose homeostasis parameters in *db/db* mice treated with BIBO3304. Four-week-old leptin receptor deficient *db/db* mice were randomized to receive placebo or oral Y1 antagonist BIBO3304 for 6 weeks. (A) Daily food intake of *db/db* mice treated with placebo or oral BIBO3304 (n = 5-6 per group). (B-C) *db/db* mice treated with placebo or BIBO3304 were fasted overnight, and intraperitoneal glucose tolerance tests (0.5 g/kg body weight) were performed. Blood glucose levels during tolerance tests were monitored and results are expressed over the time course and as area under the curve. (n = 5-6 per group). (D) Livers and white adipose tissues were isolated from 10-week-old *db/db* mice treated with placebo or BIBO3304 and Akt activation were determined. Tissues were subjected to SDS-PAGE and western blot analysis using anti-Akt phosphorylation Ser 473, and β-actin antibodies (*n* = 5-6). Results shown are a representative blot and quantitative densitometry analysis. (E-G) Ten-week-old leptin receptor deficient *db/db* mice were randomized to receive placebo or oral Y1 antagonist BIBO3304 for 6 weeks. Weekly body weights were determined. Whole body lean and fat mass as determined by EchoMRI analysis in placebo or oral BIBO3304 treated *db/db* mice (n = 4-6 per group). (H) Dissected weights of individual white adipose tissue from epididymal (Epi), inguinal (Ing) and brown adipose tissue (BAT) (n = 4-5 per group). (I) *db/db* mice treated with placebo or BIBO3304 were fasted overnight and intraperitoneal insulin tolerance tests (2.5i.u./kg body weight) were performed. Blood glucose levels during tolerance tests were monitored and results are expressed over the time course and as area under the curve. (n = 4-6 per group). (J) *Npy* and *Npy1r* mRNA expression in islets from 10-week-old *db/db* and *db/+* (n=3-10). (K-N) Pancreases from placebo or BIBO3304 treated mice were weighed and fixed in formalin and processed for immunostaining of insulin (green) and nuclear counterstained with DAPI (blue). Islet number, islet area and islet proportion were determined across two non-consecutive pancreatic sections per mouse and normalized to total pancreas section area. n = 4-6 per group. Data are means ± SEM. \**P* < 0.05, \*\**P* < 0.01; calculated by unpaired Student’s *t*-test or two-way ANOVA analysis.

